# CD8+ T cell cross-reactivity against SARS-CoV-2 conferred by other coronavirus strains and influenza virus

**DOI:** 10.1101/2020.05.20.107292

**Authors:** Chloe H. Lee, Mariana Pereira Pinho, Paul Buckley, Isaac Woodhouse, Graham Ogg, Alison Simmons, Giorgio Napolitani, Hashem Koohy

## Abstract

While individuals infected with coronavirus disease 2019 (COVID-19) manifested a broad range in susceptibility and severity to the disease, the pre-existing immune memory of related pathogens can influence the disease outcome. Here, we investigated the potential extent of T cell cross-reactivity against severe acute respiratory syndrome coronavirus 2 (SARS-CoV-2) that can be conferred by other coronaviruses and influenza virus, and generated a map of public and private predicted CD8+ T cell epitopes between coronaviruses. Moreover, to assess the potential risk of self-reactivity and/or diminished T cell response for peptides identical or highly similar to the host, we identified predicted epitopes with high sequence similarity with human proteome. Lastly, we compared predicted epitopes from coronaviruses with epitopes from influenza virus deposited in IEDB to support vaccine development against different virus strains. We believe the comprehensive *in silico* profile of private and public predicted epitopes across coronaviruses and influenza viruses will facilitate design of vaccines capable of protecting against various viral infections.

## Introduction

Faced by unprecedent health and economic crisis from the coronavirus disease 2019 (COVID-19), the scientific community is pushing forward with efforts to develop vaccines and treatments to mitigate its impact. While the severity of symptoms have been reported to be associated with age, gender and comorbidities such as cardiovascular diseases and chronic respiratory diseases^1^, the underlying mechanism of broad variation in susceptibility and severity to COVID-19 is not fully understood^2^. It is however accepted that an altered immune response is a key contributor to pathology^3,4^, and the balance between generation of protective and pathological immune responses by the host may be a vital factor governing the disease outcome.

As immune memory by related pathogens has shown to help reduce severity and spread of the diseases ^5–7^, pre-existing immunity through cross-reactivity to familial coronavirus strains may provide individuals with protection or enhanced susceptibility against the severe acute respiratory syndrome coronavirus 2 (SARS-CoV-2) without prior exposure^8–10^. Therefore, we aim to characterize the potential for the existing immune memory by other coronaviruses and influenza virus to fight against SARS-CoV-2 and further identify targets for developing a ‘universal’ vaccine against coronaviruses.

The strains infecting humans belong to either alpha and beta genera of coronavirus. The alphacoronavirus contains human coronavirus 229E (HCoV-229E) and HCoV-NL63, while the betacoronavirus contains HCoV-OC43, and HCoV-HKU1, middle east respiratory syndrome coronavirus (MERS-CoV), SARS-CoV and SARS-CoV-2^11^. It is known that NL63, 229E, OC43, and HKU1 usually cause only mild to moderate symptoms such as cough, runny nose, fever and sore throat like the common cold^12^, whereas MERS-CoV and SARS-CoV cause more severe symptoms including respiratory tract disease.

In this study, we investigated the level of T cell antigen cross-reactivity across the seven alpha and betacoronavirus strains, evaluated the risk of self-reactivity from SARS-CoV-2 predicted epitopes and identified targets for vaccine developments against coronavirus and influenza virus. We first predicted the potential of peptides to be presented by ten prevalent HLA alleles and eliciting CD8+ T cell responses, and generated a comprehensive *in silico* profile of public and private predicted epitopes. We also expanded the map of cross-reactivity from exact matching peptides to those with a high sequence similarity, resulting in addition of 264 and 283 public SARS-CoV-2 predicted epitopes by allowing one and two amino acid mismatches, respectively. Moreover, to assess the risk of self-reactivity and immunopathology, we compared SARS-CoV-2 predicted epitopes with human proteome sequences and detected ten predicted epitopes that are single amino acid variant from their counterparts in the human proteome. Lastly, to identify peptides to support development of vaccines against coronavirus and influenza virus, we compared our list of predicted epitopes from coronaviruses with epitopes from influenza virus deposited in IEDB and detected epitopes with a modest sequence similarity that are shared across multiple coronavirus strains.

## Results

### Shared predicted epitopes among coronavirus strains

To first evaluate the homology of proteome sequences among alpha and betacoronavirus strains, we conducted sequence alignment and generated a phylogenetic tree of encoded proteins between NL63, 229E, OC43, HKU1, MERS-CoV, SARS-CoV and SARS-CoV-2 (see Methods). Based on sequence alignments (example alignment of spike protein illustrated in Supplementary Figure 1) and phylogenetic trees of encoded proteins, the alpha and betacoronavirus strains shared a high sequence similarity within their own genera, i.e. higher similarity between NL63 and 229E than with betacoronaviruses. In particular, for betacoronavirus strains, phylogenetic analysis showed higher similarity between OC43 and HKU1, and between SARS-CoV and SARS-CoV-2. Notably, MERS-CoV showed relatively distinct proteome sequences to all other coronavirus strains included in this study (Figure 1A for spike protein and Supplementary Figure 2 for other encoded proteins).

**Figure 1.**
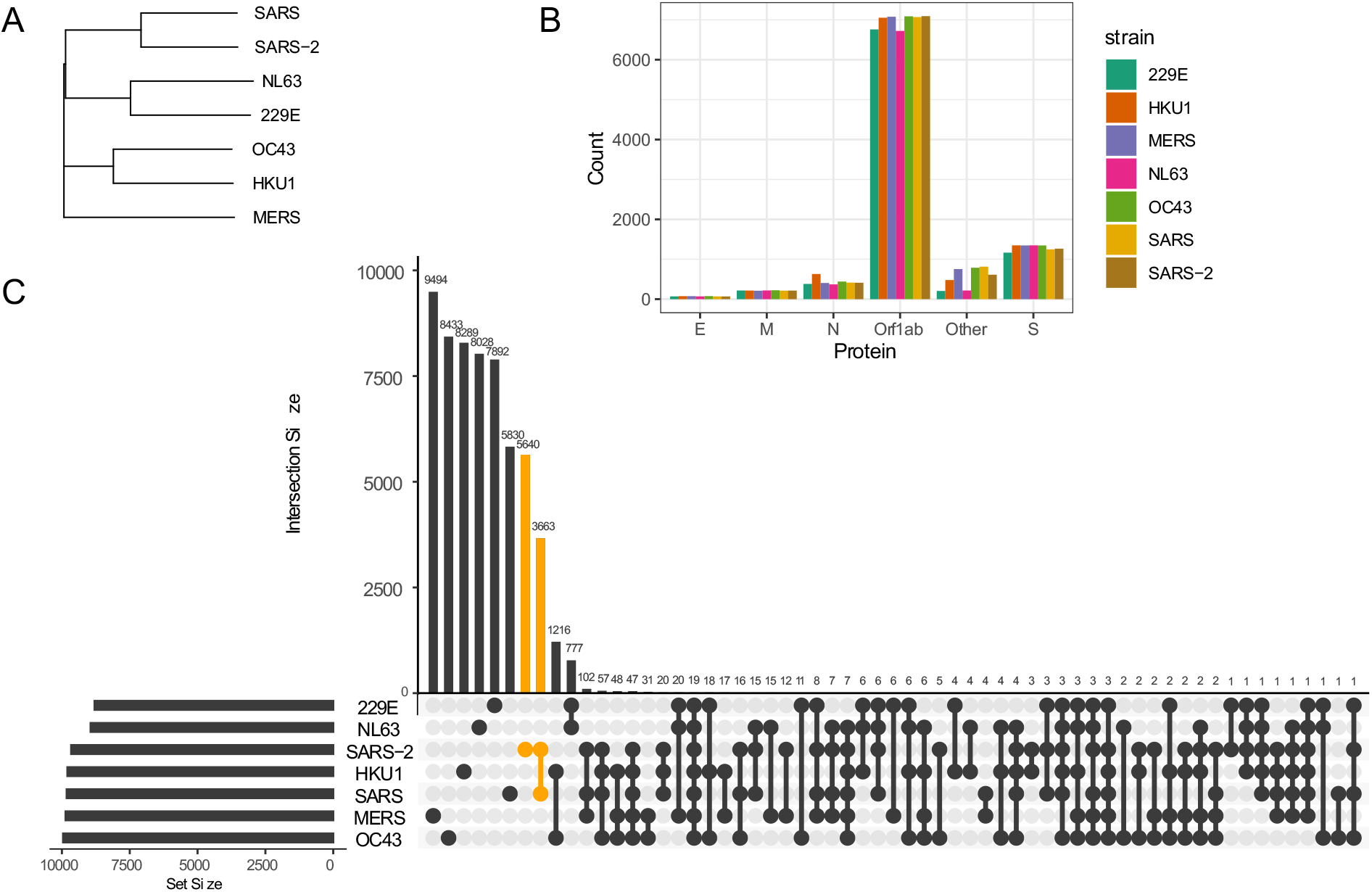
Sequence homology between coronavirus strains. A. Phylogenetic tree of spike protein sequences between NL63, 229E, OC43, HKU1, MERS-CoV, SARS-CoV and SARS-CoV-2 strains. B. Number of 9-mer peptides generated from each coronavirus strains grouped by functional proteins. C. Number of shared and private 9-mer peptides between coronavirus strains.

To identify the conserved 9-mer peptides across coronavirus strains, we first generated 9-mer peptides from encoded proteins of coronavirus strains (Figure 1B) and detected public peptides with identical matches (Figure 1C). Notably, there were 3663 shared 9-mer peptides between SARS-CoV and SARS-CoV. Given the longest open reading frame of replicase polyproteins (Orf1ab) with an average of 7002 amino acids across coronavirus strains, many public peptides were derived from Orf1ab, with 19 peptides shared across all studied strains (Figure 3 first panel).

We then investigated antigen presentation potential of 9-mer peptides for 10 most prevalent HLA alleles corresponding to MHC class I (HLA-A, HLA-B and HLA-C alleles). These include HLA-A*0101, 0201, 0301, 2402, HLA-B*0702, 4001, 0801 and HLA-C*0702, 0401, 0701. The MHC presentation was predicted by netMHCpan v4.0^13^ (see Methods). Generally, there was a relatively high number of peptides predicted to bind HLA-C alleles while HLA-B alleles had the lowest number of predicted binders (Figure 2A). We identified on average 1380 (SD = 45) peptides predicted to bind at least one HLA allele (Figure 2B), which is ~15% of total number of 9-mer peptides across different strains. Of interest, there were few peptides predicted to bind by at least 7 different HLA alleles, of which 9 peptides were derived from SARS-CoV-2. The peptides predicted to bind 8 HLA alleles and their derived proteins are listed in Supplementary Figure 3.

**Figure 2.**
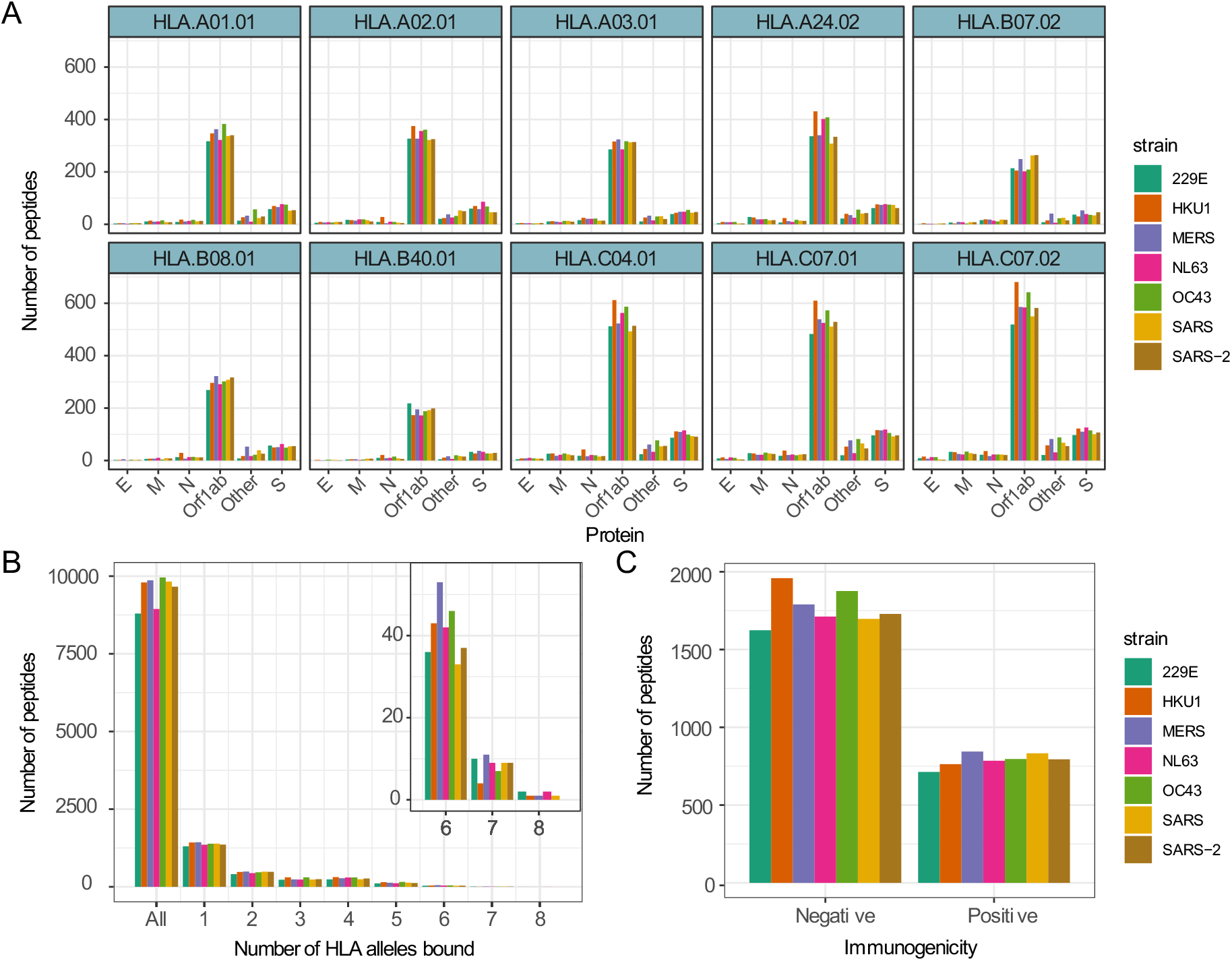
Number of 9-mer peptides predicted to bind HLA alleles and are immunogenic. A. Number of 9-mer peptides from each coronavirus strains predicted to bind annotated HLA alleles. B. Number of peptides predicted to bind equal to specified number of HLA alleles. C. Number of peptides predicted to trigger T cell response by Repitope prediction.

Although viral antigen presentation is a vital step in triggering immune responses, not all MHC binding peptides are immunogenic. We therefore set out to predict T cell immunogenicity, i.e. the ability of a peptide to elicit a T cell response, of all peptides predicted to bind at least one HLA allele. In an ongoing unpublished study, we have benchmarked the existing immunogenicity predicting models and as a result recognized a recently published model called Repitope^14^ as the best performing existing model to predict immunogenicity of viral epitopes.

We therefore utilized Repitope (see Methods) and identified in total 4894 out of 16096 (~30%) unique predicted binders to be immunogenic (subsequently referred to as predicted epitopes), and the proportion of such predicted epitopes was comparable across different strains (Figure 2C, Supplementary Figure 4). On average, we detected 429 (SD=26) epitopes predicted to bind at least one HLA type and trigger T cell response. The full list of predicted immunogenic and nonimmunogenic HLA-binders are provided in Supplementary Table 1.

With the pool of predicted epitopes, we generated Venn diagrams to illustrate private and public epitopes across different coronavirus strains (Figure 3, Supplementary Figure 5). From a total of 794 predicted epitopes from SARS-CoV-2, 411 were private while the remaining were public of which 350 were shared between SARS-CoV-2 and SARS-CoV. We detected only one predicted epitope SLAIDAYPL public across all strains. Given the long sequence of Orf1ab protein, many public epitopes were derived from Orf1ab.

**Figure 3.**
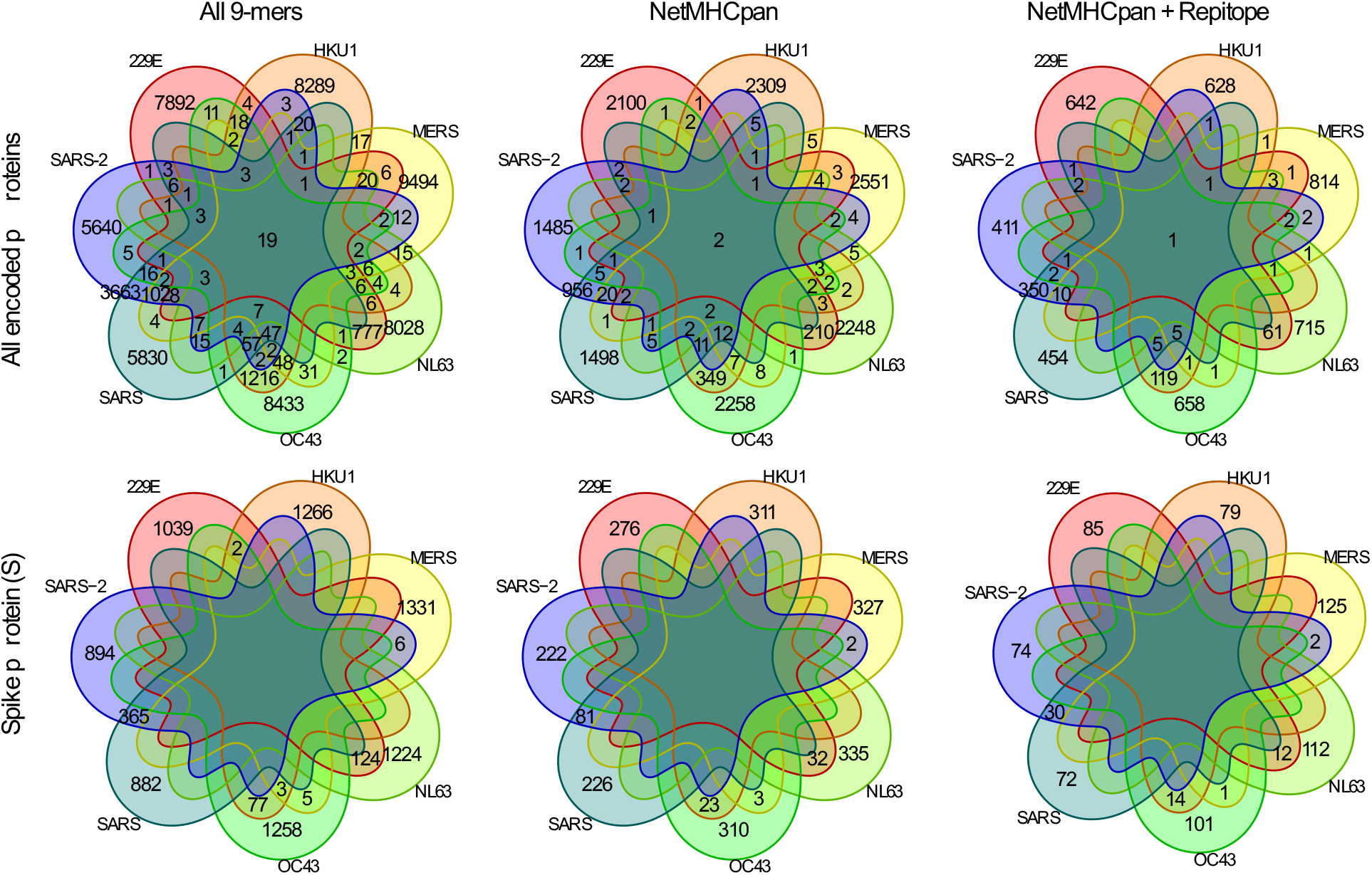
Comparing private and public 9-mer peptides from a complete set of peptides to after MHC presentation prediction by NetMHCpan and immunogenicity prediction by Repitope.

For a validation of predicted epitopes, we have compared the list of predicted epitopes with epitopes characterized by *in vitro* T cell assays and deposited in Immune Epitope Database and Analysis Resource (IEDB). We retrieved all 35,224 peptides in IEDB (as of 13-02-2020), presented on either MHC-I or MHC-II, reported positive in T cell assays and have human as the host organism, then compared with the list of 4894 unique predicted epitopes. We identified 66 unique 9-mer predicted epitopes matching with IEDB epitopes from four coronavirus strains, 229E, NL63, SARS-CoV and SARS-CoV-2 (Table 1. Supplementary Table 2). In particular, the matching epitopes from NL63 and SARS-CoV-2 were public peptides with 229E and SARS-CoV respectively. Consequently, all 229E and NL63 matching epitopes in IEDB were derived from 229E whereas all SARS-CoV and SARS-CoV-2 matching epitopes in IEDB were derived from SARS-CoV.

**Table 1.** Number of unique peptides by strain having matching pattern with epitopes deposited in IEDB

| Strain | Number of unique peptides by strain |
| --- | --- |
| 229E | 2 |
| NL63 | 1 |
| SARS | 64 |
| SARS-2 | 31 |

### Public epitopes between SARS-coV-2 and other coronavirus strains by high sequence similarity

In addition to public peptides by exact matches, predicted epitopes with a high sequence similarity may also trigger cross-reactivity across coronavirus strains. Here, we expanded the previous set of public predicted epitopes of SARS-CoV-2 to include those with up to two amino acid mismatches (Table 2, Supplementary Table 3). In addition to 629 predicted epitopes from SARS-CoV-2 shared with other coronavirus strains, there can be an increase of 264 and 283 public epitopes by allowing one or two amino acid mismatches respectively.

**Table 2.** Number of shared predicted epitopes between SARS-CoV-2 and other coronavirus strains by allowing up to two mismatches.

| Strains | Exact match | 1 mismatch | 2 mismatches |
| --- | --- | --- | --- |
| 229E | 22 | 10 | 30 |
| HKU1 | 48 | 44 | 56 |
| MERS | 57 | 44 | 35 |
| NL63 | 17 | 17 | 36 |
| OC43 | 52 | 41 | 58 |
| SARS | 433 | 108 | 68 |
| **Total** | **629** | **264** | **283** |

### Autoimmune potential of SARS-CoV-2 derived predicted epitopes

In identifying targets for vaccination, peptides identical or highly similar to those of host organism may diminish the T cell response due to exclusion of high-affinity T cells by thymic negative selection and/or may induce autoimmune response. Thus, we analyzed the sequence similarity of predicted epitopes with their best matching counterparts in human proteome to hint on their potential risks.

By comparing predicted epitopes from SARS-CoV-2 with the human proteome, none of the predicted epitopes had identical match but we detected 10 and 184 epitopes with one and two amino acid mismatches respectively with their human proteome counterparts (Figure 4A). The predicted epitopes differ by only one amino acid is listed in Table 3.

**Figure 4.**
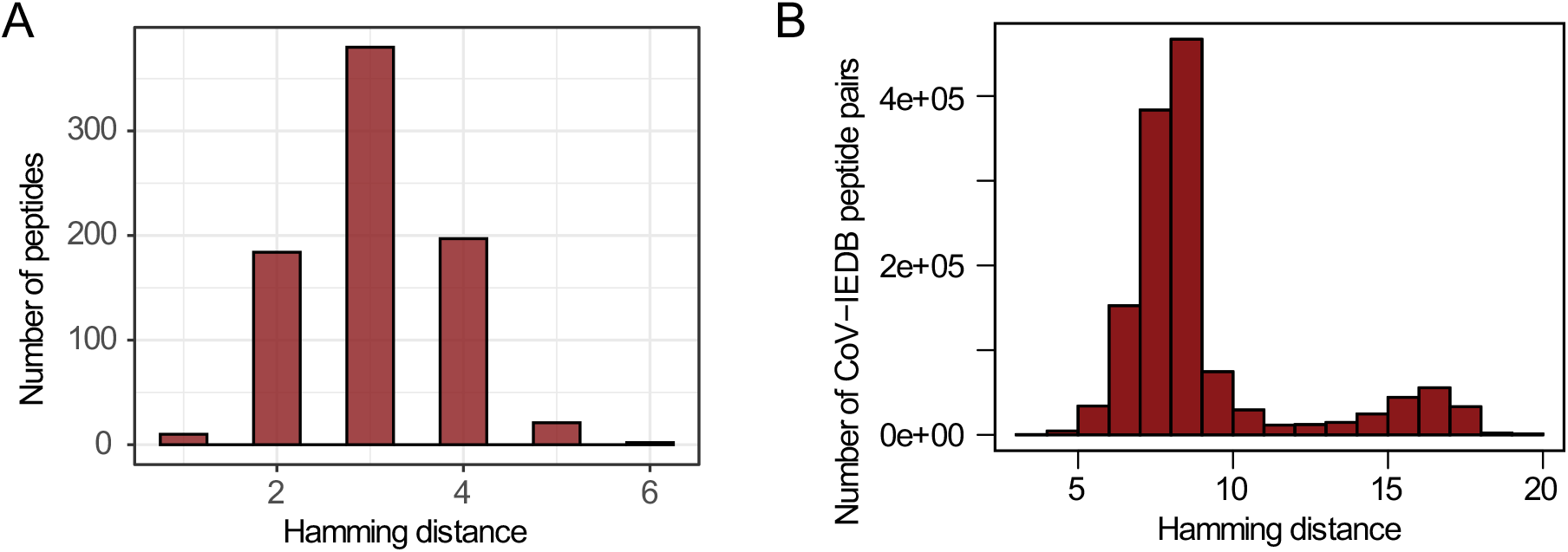
Sequence similarity with human proteome and influenza virus epitopes deposited in IEDB. A. Distribution of hamming distance between SARS-CoV-2 derived peptides and human proteome counterparts (the region most similar to corresponding virus peptides). B. Distribution of hamming distance between coronavirus derived peptides and all influenza virus epitopes deposited in IEDB.

**Table 3.** Predicted epitopes from SARS-CoV-2 that are single amino acid variants of human proteome counterparts.

| **SARS-CoV-2** | **Human proteome** |
| --- | --- |
| FLALITLAT | LLALITLAT |
| GDAALALLL | GAAALALLL |
| GLPGTILRT | GQPGTILRT |
| GLTVLPPLL | GLTVLPALL |
| IPIGAGICA | IYIGAGICA |
| IVNSVLLFL | TVNSVLLFL |
| QLSLPVLQV | QLLLPVLQV |
| SLPINVIVF | SLPINVQVF |
| TPGSGVPVV | EPGSGVPVV |
| VLPQLEQPY | VLPQNEQPY |

### Cross-reactivity between coronavirus and influenza virus

Due to the prevalence of disease caused by influenza virus and its tendency to gain mutations (antigenic drift) and reassortment among subtypes of virus (antigenic shift), there have been multiple attempts to develop generic vaccines protective against all influenza viruses to reduce the severity of infection and spread of disease^15^. Along with cross-reactivity among influenza virus strains, the cross-reactivity between coronavirus and influenza virus would benefit the community to combat recurrence of diseases caused by both strains. Here, we compared the 4894 unique predicted epitopes from all coronavirus strains with 1334 unique MHC-I influenza virus-derived epitopes (1298 influenza A virus and 36 influenza B virus) deposited in IEDB.

Due to a relative sequence dissimilarity between influenza virus and coronavirus, there were no peptides with identical match between two strains and all peptides were distinct by at least three amino acids (Figure 4B). But interestingly, among those with three amino acid differences, there were predicted epitopes shared across multiple coronavirus strains as exemplified in Table 4 (full list provided in Supplementary Figure 6). These public peptides pose a potential to be cross-protective within coronavirus strains and given that they share a modest sequence similarity with epitopes derived from influenza virus, also pose a potential to cross-react against influenza virus.

**Table 4.**
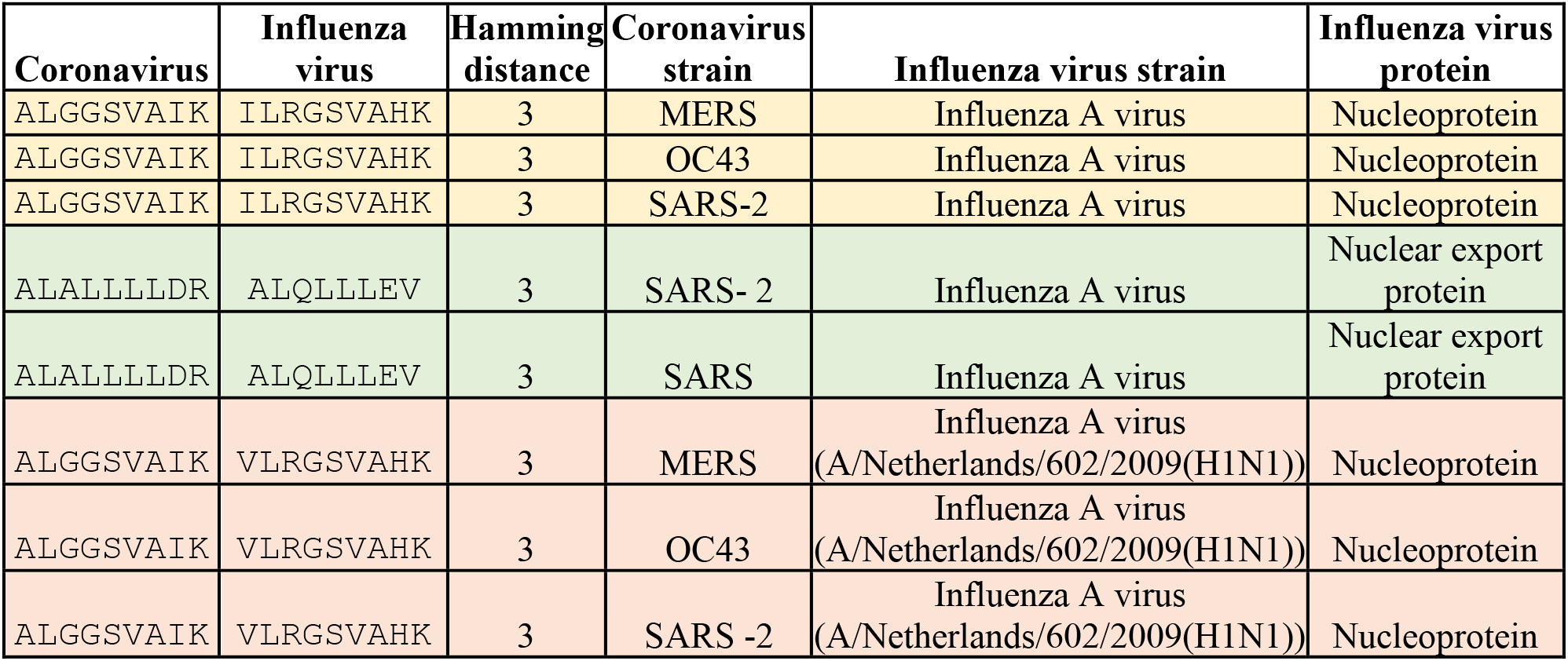
Example of public predicted epitopes from coronavirus strains with modest sequence similarity with influenza virus epitopes.

## Discussion

The severity and recurrence of coronavirus disease outbreaks pose ongoing global threat, and prompts the need for better understanding of potential cross-protection by prior infection of familial coronaviruses to mitigate the current spread and prevent future pandemics. Hereby, in continuation of our previous study to identify vaccine target for SARS-CoV-2^16^, we sought to determine the extent of antigen cross-reactivity amongst coronavirus strains.

Taking a step ahead of previous efforts to study potential immune recognition of SARS-CoV-2 by earlier infections of common coronaviruses based on MHC presentation predictions^17–21^, we i) shortlisted predicted epitopes by taking immunogenicity potential of predicted binders into account, ii) validated the prediction by comparing with epitopes deposited in IEDB, and iii) expanded the map of public predicted epitopes to accommodate up to two amino acid variants. To aid design of vaccines targeting both coronaviruses and influenza viruses, we further analyzed correlates of the predicted epitopes from coronaviruses with epitopes from influenza virus.

Along with the *in silico* profile of predicted epitopes shared across coronaviruses, we evaluated the potential risk of self-reactivity imposed by high sequence similarity with the human proteome. Considering the reports of lung, heart, liver, intestine, genital and kidney failures by autoimmune disorders in COVID-19 patients^22^, peptides from SARS-CoV-2 may carry high risk of immunopathology and should be carefully selected to proceed for vaccination.

In regards to the accuracy of predictive models utilized in this study, the algorithms to predict MHC presentation has matured significantly in the last decade by training with extensive datasets, especially for the most common HLA types. On the other hand, it is worth noting that predicting immunogenicity is challenging and not a fully solved problem. Therefore, although the best performing models have been used for classifying immunogenicity, the predictions are suboptimal and should be taken with caution.

We believe that our comprehensive profile of private and public predicted epitopes across coronaviruses and influenza virus will assists biologists with targeted function validation, and facilitate design of vaccines capable of protecting against multiple prevalent virus strains.

## Methods

### Retrieval of coronavirus proteome sequences

The proteome sequences of coronavirus strains were obtained from NCBI. The reference numbers for these sequences are NC_002645.1 (229E), NC_006577.2 (HKU1), NC_019843.3 (MERS-CoV), NC_005831.2 (NL63), NC_006213.1 (OC43), NC_004718.3 (SARS-CoV) and NC_045512.2 (SARS-CoV-2).

### Sequence alignment and phylogenetic tree of encoded proteins in coronaviruses

For multiple sequence alignment and phylogenetic tree generation, the open reading frames were grouped by their encoded proteins – spike protein (S), envelope protein (E), membrane protein (M), nucleocapsid protein (N), replicase polyprotein (Orf1ab) and other encoded regions (Other) – prior to analysis. The multiple sequence alignment was conducted using ‘msa’ function from R msa v1.4.3 package and visualized by ‘msaPrettyPrint’ function. The phylogenetic tree was produced by plotting ‘identity’ distance generated by ‘dist.alignment’ function from R seqinr v3.6.1 package.

### Peptide generation from proteome sequences

Each encoded proteins were fragmented into 9-mer peptides by scanning the proteome with a window of 9 amino acids and step length of 1 amino acid. For strains containing two open reading frames annotated with the same functional protein e.g. 229E, MERS-CoV, SARS-CoV and SARS-CoV-2 having two open reading frames annotated for Orf1ab, 9-mer peptides were generated from both encoded proteins and unique set of peptides were selected for subsequent analysis.

### MHC presentation prediction

The antigen presentation of MHC was predicted using NetMHCpan v4.0^13^, a model trained on binding affinity and eluted ligand data, against HLA-A*0101, 0201, 0301, 2402, HLA-B*0702, 4001, 0801 and HLA-C*0702, 0401, 0701 alleles. Peptides with rank score <= 2.0 were categorized as positive HLA-binder.

### Immunogenicity Prediction

The immunogenicity potential was predicted by R package Repitope for peptides that passed netMHCpan filtering i.e those predicted to bind at least one HLA allele. The Repitope utilizes amino acid descriptors and TCR-peptide contact potential profiling (CPP)-based features to label immunogenicity. The Repitope package was retrieved from GitHub repository (https://github.com/masato-ogishi/Repitope.git).

After feature computation and feature selection, we utilized the published Repitope ‘MHCI_Human’ model to extrapolate probabilistic immunogenicity scores for our dataset and made binary classification of immunogenicity. This classification was based on a threshold computed from the ROC curves of the MHCI_Human immunogenicity prediction model and was calculated using the youden index that maximizes *1-sensitivity+specificity*. Probabilistic scores were extracted from the original model for a subset of peptides that were identical to peptides in the model’s training dataset.

### Visualization of private and public peptides

The conservation of peptides across coronavirus strains before and after MHC binding and immunogenicity prediction were visualized by ‘venn’ function from R venn v1.9 package and upset function from R UpSetR v1.4.0 package. To identify shared peptides with up to two amino acid tolerance, the best matching peptides were identified by ‘pairwiseAlignment’ from Biostrings v2.40.2 package using BLOSUM62 matrix, gapOpening of 100 and gapExtension of 100, followed by hamming distance to filter only peptide pairs with less than or equal to two amino acid difference.

### Sequence similarity with human proteome and epitopes deposited in IEDB

The sequence similarity of peptides derived from SARS-CoV-2 with human proteome counterparts was computed by first identifying best global-local alignment by ‘pairwiseAlignment’ function from Biostrings v2.40.2 package with a high gap penalty, gapOpening of 100 and gapExtension of 100. The number of mismatches between best aligned pair was computed by hamming distance using ‘stringdist’ function from R stringdist v0.9.5.5 package. Similarly, the sequence similarity between coronavirus peptides and epitopes from IEDB was compared by global-local alignment following by computing hamming distance to find the best matching pairs.

## Author contributions

H.K. and G.N. conceived and designed the study. C.H.L conceived and designed computational approaches, conducted all data analysis and generated figures and tables. P.B. conducted immunogenicity prediction. I.W. conducted MHC presentation prediction. C.H.L, M.P.P, G.O., A.S., H.K., and G.N. wrote the manuscript, and I.W. and P.B. contributed to methods. A.S. and H.K. supervised the project. All authors discussed the results and contributed to the final manuscript.

## Acknowledgements

This work has been supported by Medical Research Council. H.K. is funded by MRC HIU core funding and MC_UU_00008. C.H.L is funded by UK National Institute of Health Research (NIHR). A.S. is funded by a Wellcome Investigator Award (219523/Z/19/Z), the UK Medical Research Council, NIHR, awards from Bristol-Myers Squibb and UCB. A.S. is an NIHR Senior Investigator and acknowledges support from the Oxford NIHR Biomedical Research Centre. The views expressed are those of the author(s) and not necessarily those of the NHS, The NIHR, or the Department of Health, UK. PB and IW are funded by MRC grants MC_UU_00008 and A93261 respectively. M.P.P. and G.N. are funded by the department of Health and Social Care (project number:16/107/01) as part of the UK Vaccine Network.

## Declaration of interest

G.O. has served on advisory boards or holds consultancies or equity with Eli Lilly, Novartis, Janssen, Sanofi, Orbit Discovery and UCB Pharma, and has undertaken clinical trials with Atopix, Regeneron/Sanofi, Roche.

## Supplementary Figures

**Supplementary Figure 1.**
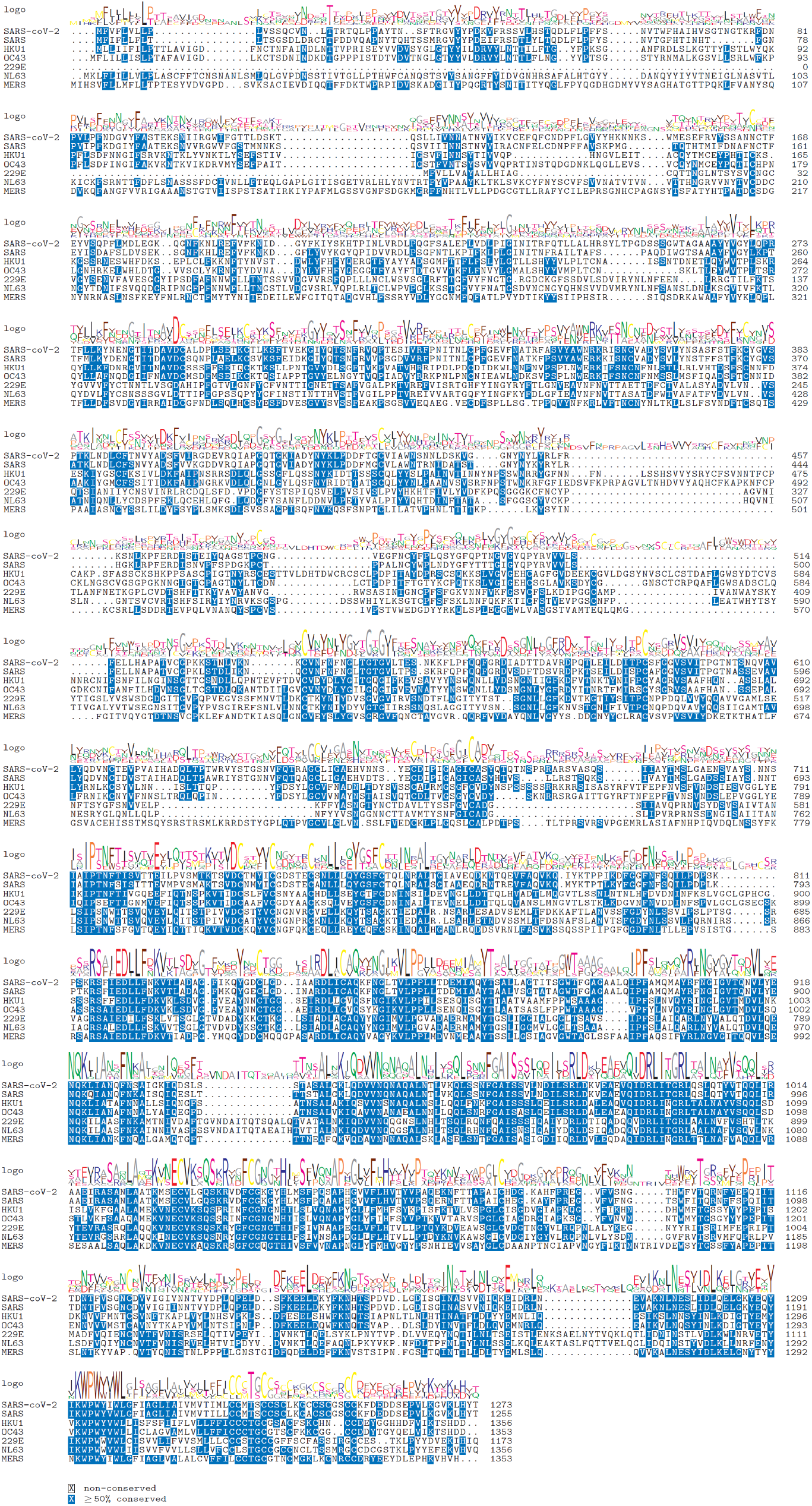
Alignment of proteome sequences of spike proteome among different coronavirus strains

**Supplementary Figure 2.**
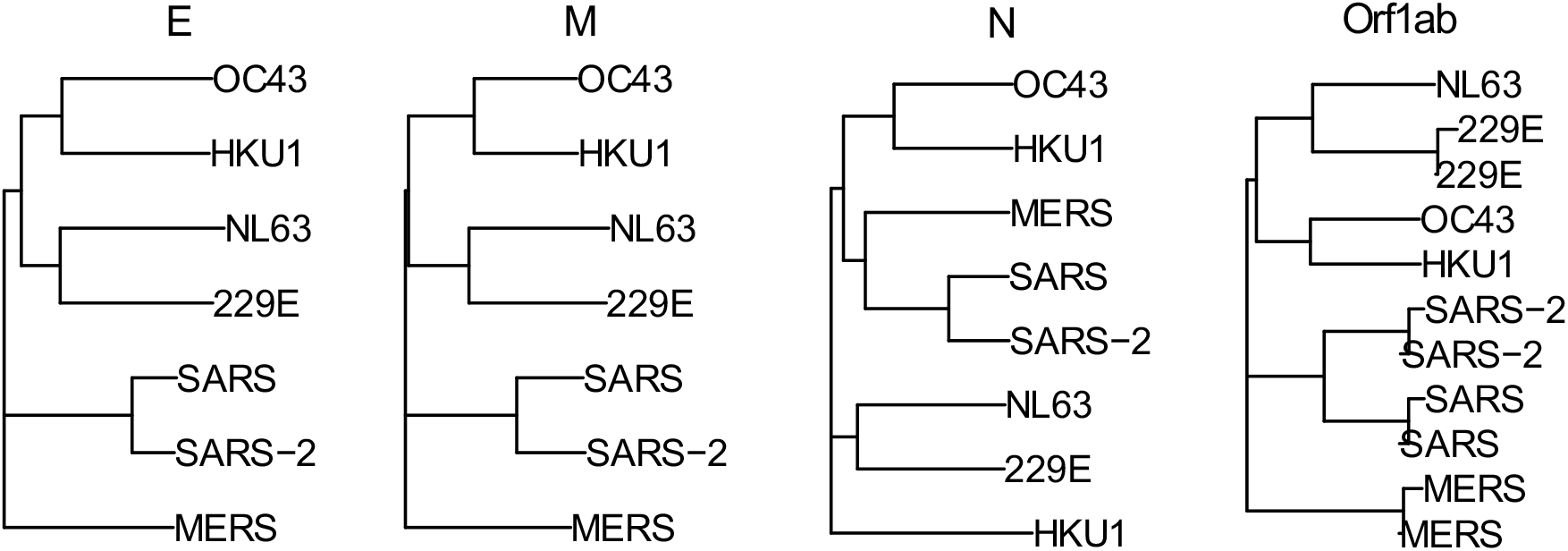
Phylogenetic tree of encoded proteins, envelope protein (E), membrane protein (M), nucleocapsid protein (N) and replicase polyprotein (Orf1ab) for seven coronavirus strains. The recurrent strain annotations are due to presence of two open reading frames annotated with the same functional protein.

**Supplementary Figure 3.**
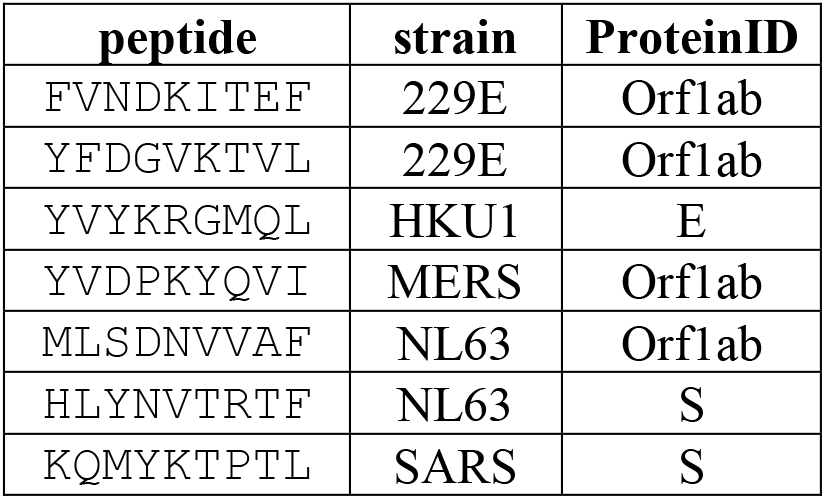
List of peptides predicted to bind 8 HLA alleles by NetMHCpan 4.0.

**Supplementary Figure 4.**
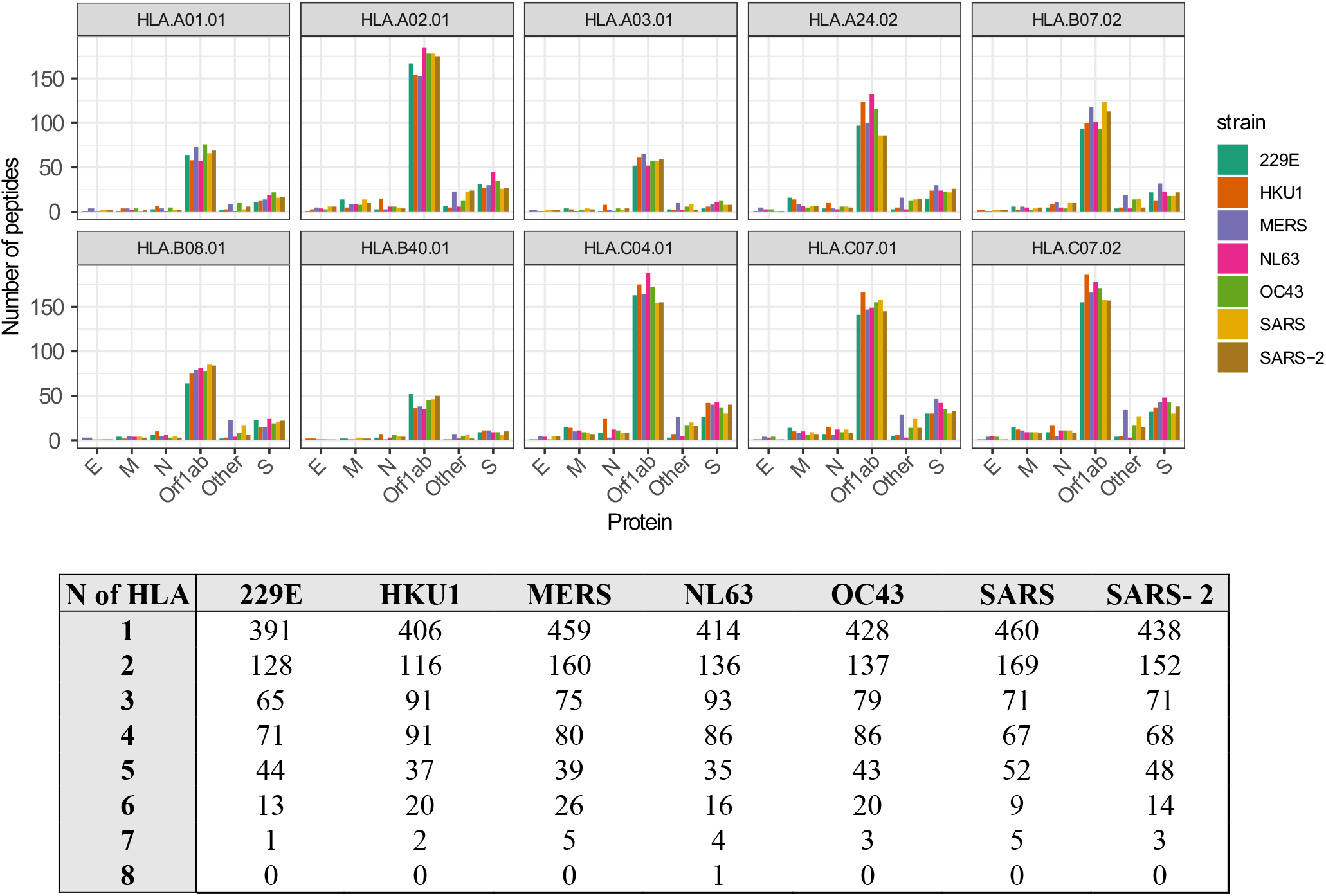
(Top) Number of peptides predicted to be immunogenic by Repitope, facet by HLA alleles predicted to bind via NetMHCpan. (Bottom) Number of predicted epitopes predicted to be N number of HLA alleles for each coronavirus strain.

**Supplementary Figure 5.**
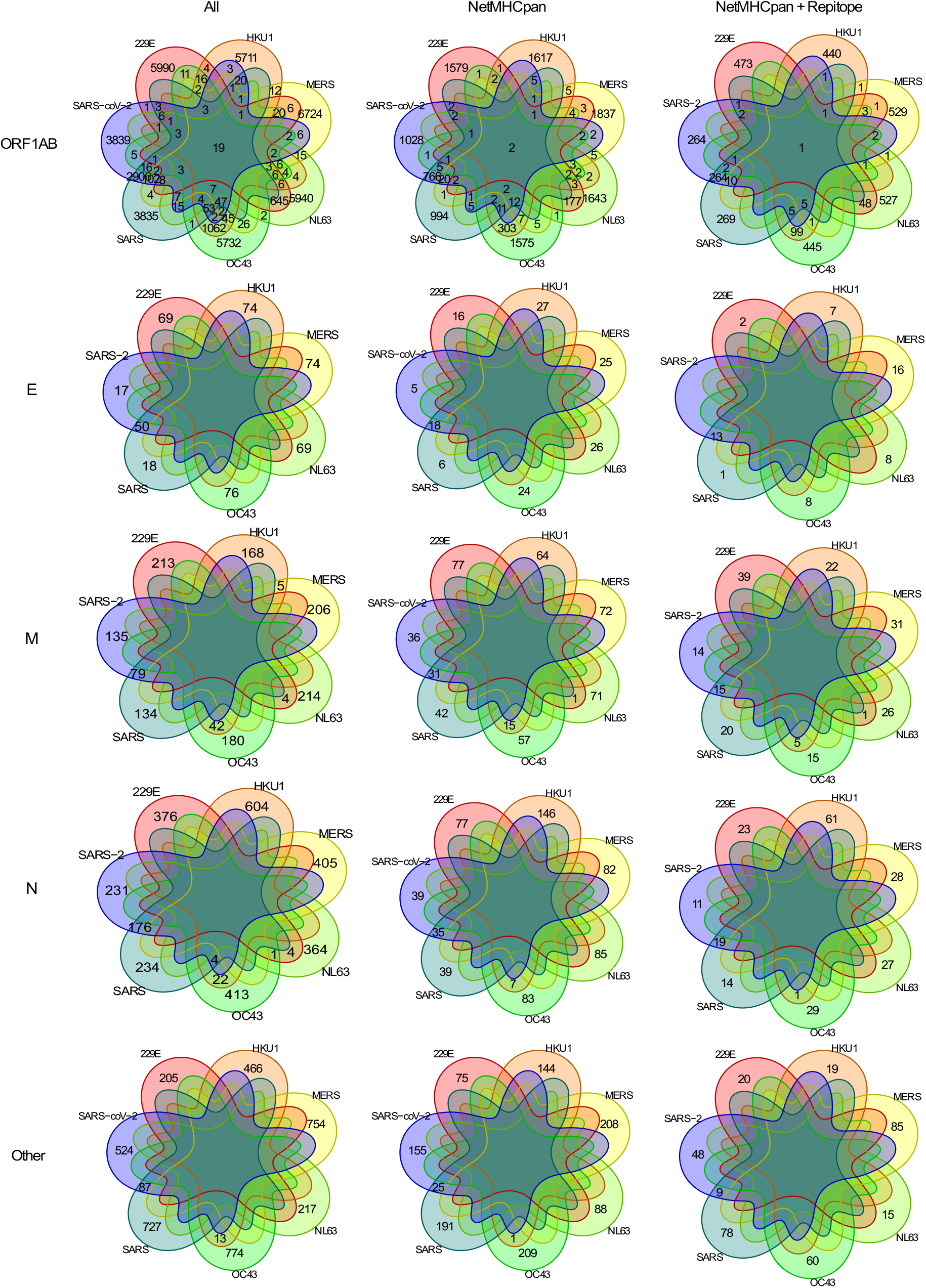
Venn diagram illustrating private and public 9-mer peptides from a complete set of peptides to after MHC presentation prediction by NetMHCpan and immunogenicity prediction by Repitope. The encoded proteins are replicase polyprotein (Orf1ab), nucleocapsid protein (N), envelop protein (E), membrane protein (M) and remaining encoded proteins combined (Other).

**Supplementary Figure 6.**
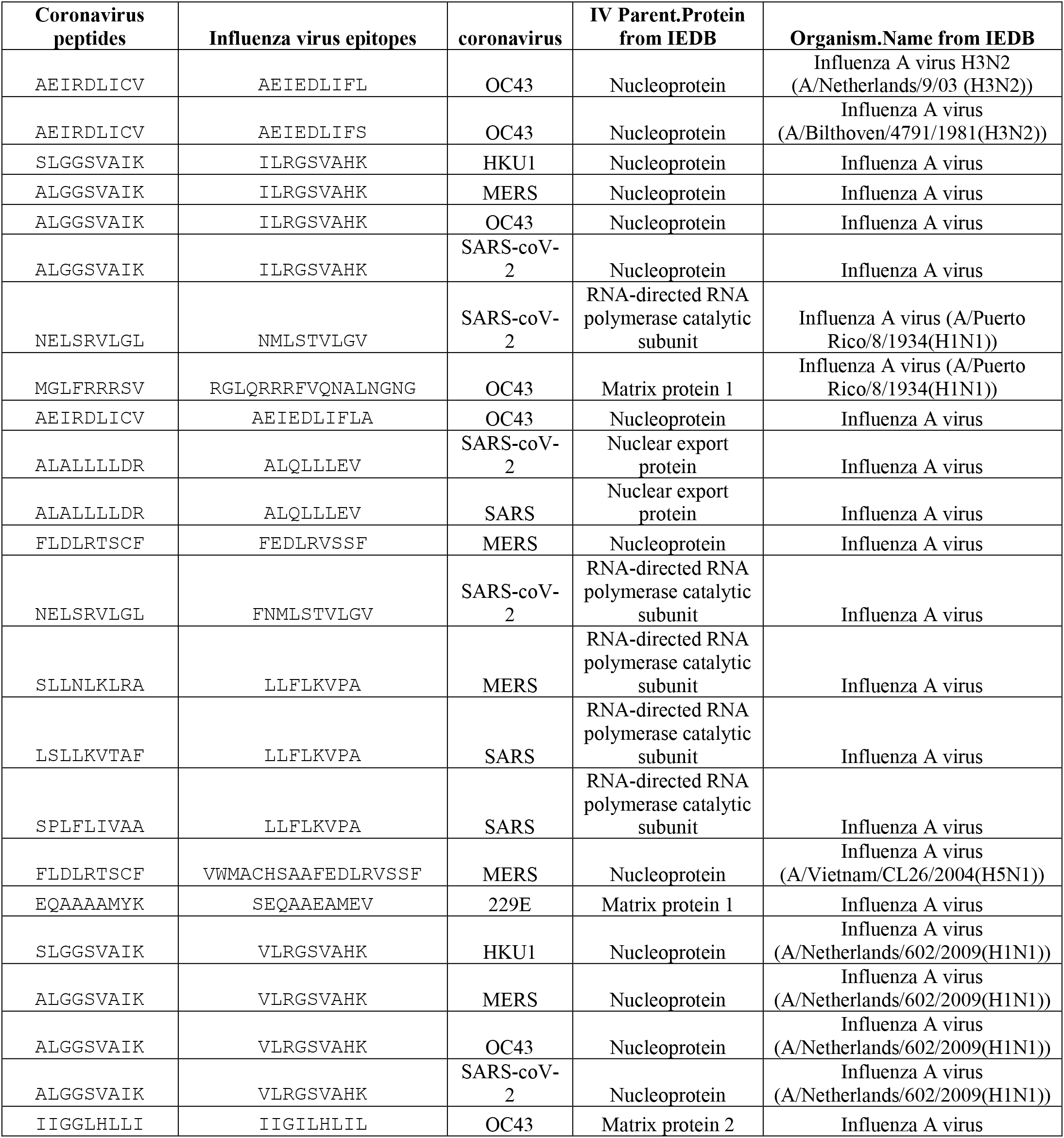
List of predicted epitopes from coronavirus and epitopes from influenza virus deposited in IEDB with a modest sequence similarity.

